# NetMD: Unsupervised Synchronization of Molecular Dynamics Trajectories via Graph Embedding and Time Warping

**DOI:** 10.1101/2025.08.12.669871

**Authors:** Manuel Mangoni, Salvatore Daniele Bianco, Francesco Petrizzelli, Michele Pieroni, Pietro Hiram Guzzi, Viviana Caputo, Tommaso Biagini, Tommaso Mazza

**Affiliations:** Department of Experimental Medicine, Sapienza University of Rome, Rome, Italy; UOS Computational Biology and Bioinformatics, Fondazione Policlinico Universitario A. Gemelli, IRCCS, Rome, Italy; Bioinformatics Laboratory, IRCCS Casa Sollievo della Sofferenza, S. Giovanni Rotondo, Italy; Department of Biochemical Sciences A. Rossi Fanelli, Sapienza University of Rome, Rome, Italy; Department of Surgical and Medical Sciences, University Magna Græcia of Catanzaro, Catanzaro, Italy

**Keywords:** Molecular Dynamics Trajectory Alignment, Graph Embedding, Dynamic Time Warping, GLUT1 deficiency syndrome

## Abstract

Molecular dynamics (MD) simulations yield detailed atomistic views of biomolecular processes, yet comparing independent trajectories is hindered by stochastic divergence. Here, we introduce NetMD, a computational approach that synchronizes and analyzes MD trajectories by combining graph-based representations with dynamic time warping. Frames are transformed into residue–contact graphs, entropy-filtered to retain variable interactions, and embedded as low-dimensional vectors. NetMD then uses time-warping barycenter averaging to align these vector trajectories, yielding a consensus “average” trajectory while pruning the outlier simulations. Applied to diverse systems, such as transporters, demethylases, and protein complexes, NetMD revealed shared multiphase dynamics and pinpointed mutation- or ligand-specific deviations. Thus, this method enables an unsupervised, time-resolved comparison of MD ensembles across conditions. It is robust, broadly applicable, and available as an open-source software, offering a powerful tool for uncovering common patterns and critical divergences in biomolecular dynamics.

## MAIN

Molecular dynamics (MD) simulations have significantly advanced our understanding of atomic and molecular behavior^1^, enabling comprehensive exploration of the structural properties and dynamic mechanisms of complex systems across biology, chemistry, and materials science. In biology, MD is a foundational tool in computational genomics, elucidating the structural implications of genetic variants in molecular frameworks, thereby enhancing our understanding of genetic disorders and providing critical insights for research and diagnostic applications^2,3^. In chemistry, ab initio MD, such as the Car–Parrinello method, makes it possible to simulate complex bond-breaking and bond-forming reactions and explore reactive potential energy surfaces effectively^44^. MD has been widely applied in materials science to the study of Ti–Al-based alloys, providing insights into microstructure changes, alloy behavior, and interface phenomena^45^, as well as probing atomic-scale phenomena in materials, such as phase-change behavior for energy storage^46^, and guiding the design and characterization of novel nanomaterials^47^. Across these disciplines, the ability to compare multiple MD trajectories is central to identifying conserved mechanisms, quantifying variability, and distinguishing condition-specific behaviors. This is as true for detecting subtle differences in mutant protein folding as it is for evaluating alternative catalytic intermediates or benchmarking structural stability in engineered materials. However, comparing independent trajectories remains challenging because stochastic divergence can obscure genuine mechanistic differences. Misaligned simulations may mask shared dynamic motifs or exaggerate differences, limiting the reliability of conclusions drawn from them. Robust trajectory comparison is therefore a critical step in deriving consistent, scientifically meaningful interpretations from MD data, regardless of whether the target system is a membrane transporter, a catalytic complex, or a nanostructured alloy.

Traditional methods for analyzing MD trajectories often rely on root-mean-square deviation (RMSD) calculations, quantitative comparisons of structural ensembles, and clustering techniques^4–7^ to compare different conformations. Although these approaches provide valuable structural comparisons, they may fail to address temporal misalignments. Methods, such as those based on stochastic synchronization^8,9^, can address the alignment of trajectories within similar systems or configurations. These methods depend heavily on shared noise sequences or initial conditions, features often absent in distinct molecular systems with divergent structures and dynamics. As such, they are ill-suited for identifying points in time where trajectories converge, or “meet,” to reveal shared patterns or critical events^10^. In fact, temporal synchronization is not merely about aligning trajectories but requires uncovering consensus points of behavior that may arise from different initial states and pathways. Recent advancements in this field include classification of ligand-unbinding pathways. In ^11^, the authors used the dynamic time-warping method to effectively align the MD trajectories. However, this technique is highly computationally demanding for high-dimensional datasets, limiting its scalability to larger molecular systems or long simulation timescales. The problem still lies with other types of dynamics, for example, the conformational changes between the open-closed states of membrane proteins, and is further compounded when dealing with mutant proteins, whose dynamics are unknown in advance, because variations in sequence and structure may lead to significant differences in folding pathways, transition states, and interaction dynamics. Membrane proteins frequently undergo asynchronous multistep transitions that combine small helical shifts with large loop rearrangements. Such complexity makes the global RMSD or simple contact-based comparisons particularly insensitive to the functional motions involved.

Here, we present NetMD, a generalizable method for MD trajectory synchronization. NetMD converts trajectory frames into residue-contact graphs, filters edges based on entropy to highlight edges that vary significantly and embeds temporally ordered graphs into a time series of low-dimensional vectors using Graph2Vec^12^. The model can also distinguish between simulations and reconstruct their temporal progressions. These embeddings are then synchronized using Dynamic Time Warping (DTW) barycenter averaging^13^, yielding a consensus trajectory. Outlier replicas that deviate from it are pruned based on the length-normalized DTW distance. Finally, hierarchical clustering on aligned embeddings assesses replica consistency, distinguishes dynamic signatures across experimental conditions (**Fig. 1**), and yields a final reference trajectory to be input for downstream analyses, such as change-point detection and deep contact inspection. Change-point detection identifies time points whenever the mean distance from the consensus replica changes. Deep contact inspection examines the persistence of residue–residue contacts during the simulation (see **Methods**).

**Fig. 1.**
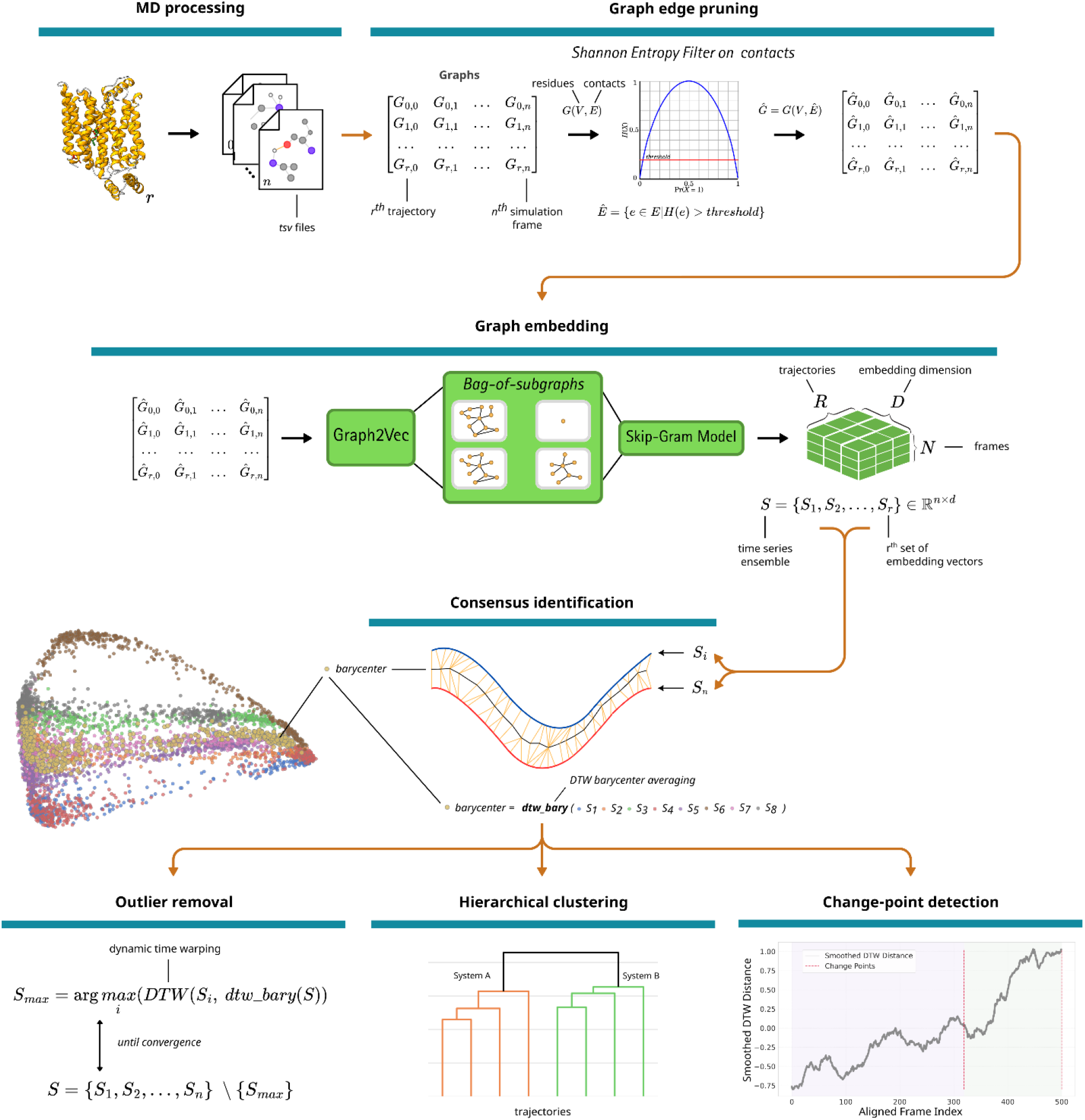
NetMD workflow. MD frames are converted into residue-contact graphs with entropy-filtered edges, embedded via a 16-dimensional Graph2Vec model into vectors, and aligned across replicas using DTW barycenter averaging (normalized by series length). Divergent trajectories were pruned, and the clustered frames yielded a state-transition network. Embeddings from the same system or under different conditions were compared using Ward’s hierarchical clustering to identify system-specific dynamic signatures. Change point detection was performed on the best replicas per system to highlight diverging states.

## RESULTS

We applied NetMD to four representative systems to demonstrate its ability to synchronize and compare trajectories across diverse molecular contexts. For human glucose transporter 1 (GLUT1), variants of which are associated with Glucose Transporter Type 1 Deficiency Syndrome, a neurological disorder of energy metabolism, we assessed NetMD’s capacity to synchronize *wild-type* and mutant trajectories, discriminate their dynamic behaviors, and determine the time windows of mutant divergence (**Fig. 2**).

**Fig. 2.**
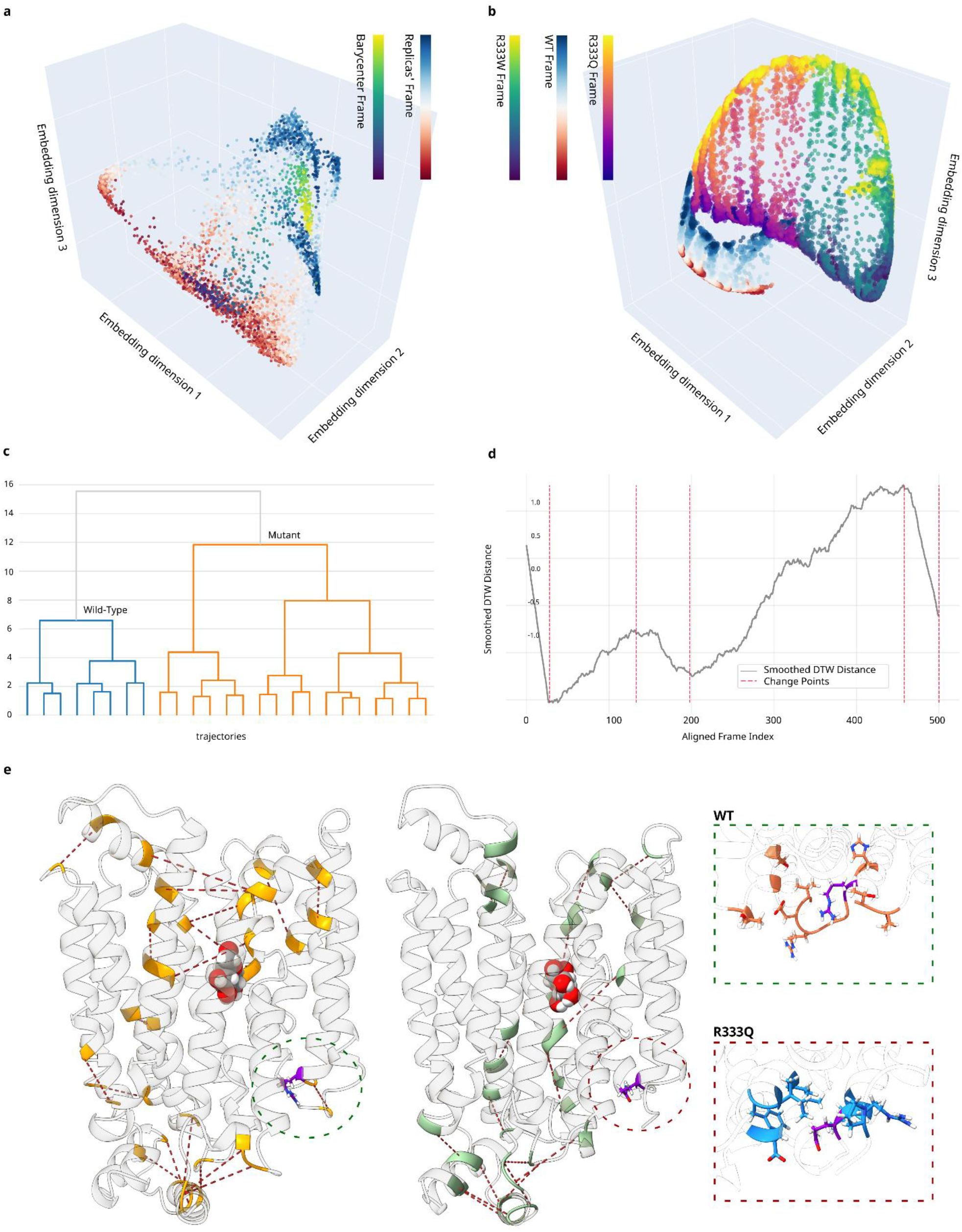
GLUT1 trajectory alignment and divergence via NetMD. (a) Three-dimensional embedding of *wild-type* GLUT1 transporter replicas overlaid with DTW barycenter trajectory (viridis). Each marker represents a single frame within the embedding space. (b) Combined 3D embeddings of the *wild-type* (RdBu) and Arg333 variants, R333Q (plasma) and R333W (viridis), showing three distinct, non-overlapping clusters that reflect their divergent conformational dynamics. (c) Final hierarchical clustering dendrogram computed on the combined embeddings, illustrating the two major branches corresponding to the *wild-type* and R333Q-R333W systems. (d) Change-point analysis of the aligned DTW distance time series with vertical dashed lines marking transition shifts. (e) *Wild-type* versus mutant favored interactions observed during the change-point window, highlighted in 3D structures as dotted lines between the two interacting residues. On the left, top-favored *wild-type* contacts are shown in yellow; on the right, top-favored mutant interactions are shown in light-green. The two panels on the far-right highlight contacts involving the mutant site: those more frequent in the *wild-type* (orange, top), and those more frequent in the mutant (blue, bottom). The mutant site was highlighted in violet. A list of the highlighted contacts is provided in **Supplementary Table 1**.

Replicate trajectories for the GLUT1 systems varied modestly in length, most extended to ~500 frames, whereas a few concluded up to 20 frames earlier. Before embedding, we applied an ensemble-specific Shannon–entropy filter to each replica of systems, i.e., the *wild-type*, R333Q, and R333W, pruning low-variability, non-informative contacts while preserving the most characteristic interactions of each ensemble. NetMD successfully synchronized and overlaid the time-series embeddings from both the *wild-type* and the two Arg333 mutant simulations, enabling unsupervised k-means clustering to reveal a reproducible three-stage glucose transport cycle, comprising an initial intake phase in which glucose approaches and binds from the extracellular side, a central conformational transition driving the outward-occluded to inward-open switch, and a final release phase where glucose exits toward the cytoplasm, directly from the aligned low-dimensional representations. This segmentation emerged consistently across GLUT1 *wild-type* and R333 mutants, confirming that NetMD can identify key mechanistic steps without prior structural labeling. Although all replicas conformed to the same global transition pattern, each showed its own subtle biologically meaningful variations in the timing and intensity of individual dynamic phases, highlighting the sensitivity of the method in capturing both the fundamental shared mechanism and divergent behaviors across simulations.

When projected using PCA followed by Spectral Embedding, the resulting embedding revealed not only the temporal progression of each replica, but also a remarkable consistency across simulations of the same system. The replicas were consistently aligned along common trajectories in the embedding space, highlighting that the essential dynamics of the systems were captured rather than random fluctuations or simulation noise. These trajectories exhibited smooth frame-to-frame progression and strong coherence across replicas, emphasizing the robustness of the method in encoding convergent molecular behavior. In each case, the replicas formed tightly clustered trajectories following alignment; however, the mutants exhibited greater deviation. This increased variation may reflect true alterations in biophysical dynamics. Alternatively, this could result from the fact that mutant trajectories were supervised to realize the three conformational states common to the *wild-type* system. However, the mutants may partially or completely disrupt these states. In such cases, the simulation may struggle to reconcile the actual mutant-induced changes with the trajectory that it is “guided” to follow, resulting in irregular behaviors or alignment inconsistencies.

The DTW barycenter traced a smooth trajectory through these states (**Fig. 2a**), whereas pruning removed the replicas that failed to closely follow the consensus. The normalized DTW distances among *wild-type* replicas spanned from approximately 0.63 (indicating near-perfect alignment) to over 1.14, reflecting the runs that deviated from the barycenter. The barycenter computed for the *wild-type* simulations showed a compact and smooth progression, suggesting a well-defined and reproducible sequence of conformational transitions. This concise trajectory underscores the strong agreement among *wild-type* replicas, reflecting the stability and convergence of the native dynamics of the system.

Hierarchical clustering distinguished *wild-type* from R333 variants coherently with embedding visualization (**Fig. 2b**), consistent with the impaired transport in these mutants (**Fig. 2c**). Applying an efficient change-point detection algorithm to the DTW-aligned distance series revealed precise intervals during which the mutant trajectories began to depart from the *wild-type* consensus; intervals that map directly onto the critical conformational steps of glucose occlusion, transition, and release (**Fig. 2d**). Notably, when we examined the frequency of residue–residue contacts within the divergence windows, we observed that the mutants lost several interactions at and around the R333 site present in the *wild-type*, while simultaneously forming new, aberrant contacts that could disrupt the normal gating mechanism (**Fig. 2e**).

Lysine-specific demethylase 6A (KDM6A) is a globular protein that catalyzes the demethylation of tri/dimethylated histone H3, which is involved in Kabuki syndrome and various cancers, including bladder cancer, pancreatic cancer, and others^14,15^.

All KDM6A GaMD replicas comprised exactly 2 500 frames each; given the initially dense contact graphs for KDM6A—with many edges present across most frames and replicas—we raised the Shannon entropy cutoff to 0.3 to selectively prune any ubiquitous and invariant interaction. This adjustment reduced the ensemble bias by removing largely constant contacts while preserving the more informative, mutation‐sensitive interactions that distinguished the *wild-type* and mutant simulations.

NetMD alignment of *wild-type* and a selected Kabuki syndrome-associated variant trajectories revealed a common core pathway in the *wild-type* trajectories and a broader embedding dispersion in the mutant (**Fig. 3a,b**). The normalized DTW distances from the barycenter among *wild-type* replicas ranged from approximately 0.82 for the closest runs up to 1.36 for the most divergent, underlining both highly representative trajectories and clear outliers, whereas in the mutant ensemble, distances spanned from roughly 0.84 up to 1.45, further underscoring the greater heterogeneity introduced by the variant. We further quantified this by computing the full pairwise normalized DTW distance matrix, which showed that *wild-type* replicates clustered tightly (average inter-replica distance ~ 2.16), whereas distances involving the R1255W variant spanned a much wider range (~2.15 up to 4.06, mean ~3.06), confirming increased heterogeneity in the mutant ensemble. Moreover, the average cross-group distance between *wild-type* and mutant replicas was ~3.5, which was substantially higher than either the *wild-type* or mutant self-distances, underscoring the clear separation between the two dynamic ensembles.

**Fig. 3.**
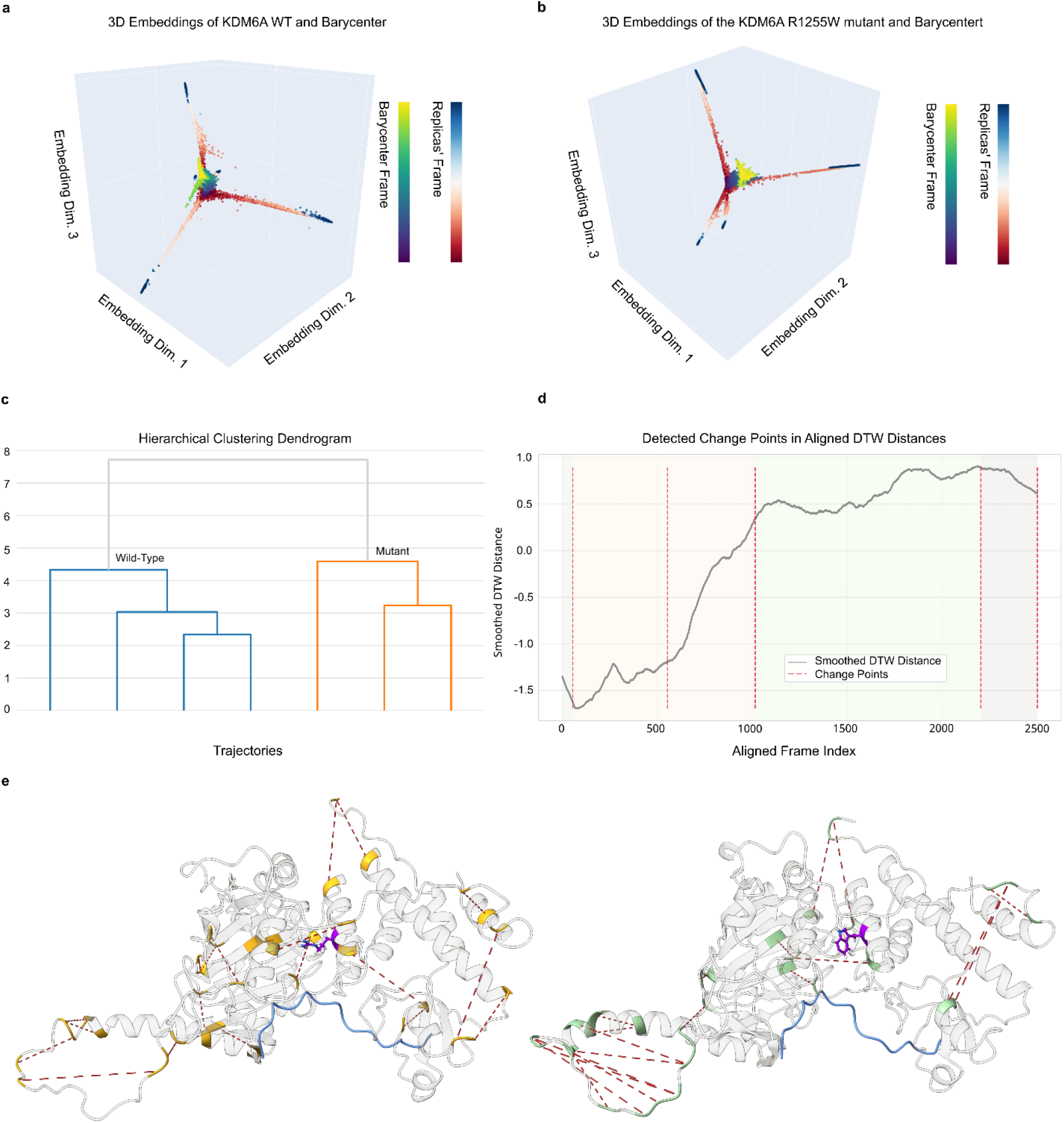
(a) Three-dimensional embedding of *wild-type* KDM6A, with the DTW barycenter trajectory overlaid (viridis). Each marker represents a single frame within the embedding space. Differences in embedding space represent the unsupervised nature of the simulations; (b) Three-dimensional embedding of the mutant KDM6A, with the DTW barycenter trajectory overlaid (viridis). (c) Final hierarchical clustering dendrogram computed on the combined embeddings, illustrating the two major branches corresponding to *wild-type* and the mutant. (d) Change-point analysis of the aligned DTW distance time series, with vertical dashed lines marking transition shifts. (e) *Wild-type* versus mutant favored interactions observed during the change-point window, highlighted in 3D structures as dotted lines between the two interacting residues. On the left, top-favored *wild-type* contacts are shown in yellow; on the right, top-favored mutant interactions are shown in light-green. The mutant site was highlighted in violet. A list of the highlighted contacts is provided in **Supplementary Table 2**.

In the case of KDM6A, embeddings from the mutant grouped into clusters that were clearly separate from those of the *wild-type* simulations, faithfully reflecting the known perturbation of histone H3 binding and the resulting alterations in domain motion (**Fig. 3c**). Change-point detection highlighted the intervals at which mutant and *wild-type* trajectories diverged (**Fig. 3d**). We focused our contact analysis on change-point intervals, where the distances exhibited pronounced increases. Within these windows, even without a predefined target event, we observed significant contact-frequency discrepancies at the mutation site and its immediate surroundings, highlighting the sensitivity of the methodology to subtle dynamic variations (**Fig. 3e**). This application illustrates NetMD’s ability to align *wild-type* and variant trajectories and pinpoint divergence windows in longer, nontargeted simulations.

We further demonstrated NetMD’s applicability to large assemblies by analyzing coarse-grained MD trajectories of human mitochondrial Complex I, where each bead represents a group of atoms, and simplified backbone contact maps were sufficient to drive the synchronization and clustering process.

Each coarse-grained Complex I trajectory consisted of 5715 frames. Each Complex I ensemble—*wild-type*, single mutant, and triple mutant —was subjected to its own Shannon–entropy filter, removing uniformly persistent backbone contacts and highlighting network changes driven by the specific ND6 mutations around TMH3. Sparser contact graphs constructed using only backbone interactions within a 15 Å radius around the ND6 mutation sites captured the essential conformational transitions of TMH3 and showed distinct clustering of the *wild-type*, single, and triple mutants. Embeddings reflected increasing divergence with mutation burden in both the embedding space and dendrogram (**Supplementary Data 1**), particularly for the triple mutant known to severely impair function, which is consistent with functional severity^16^.

To quantitatively assess alignment quality, we computed the full pairwise normalized DTW distance matrix: in the *wild-type* ensemble, distances between replicates ranged from 1.09 to 1.38 (mean ≈ 1.20), indicating tight coherence; single-mutant distances spanned 1.07–1.22 (mean ≈ 1.15) and triple-mutant distances rose to 1.43–1.59 (mean ≈ 1.49), demonstrating substantially greater heterogeneity. Cross-group distances averaged ~2.19 (*wild-type* vs. single), ~2.42 (*wild-type* vs. triple), and ~2.31 (single vs. triple), with the triple-mutant replicates lying furthest from the *wild-type* ensemble, reflecting their more pronounced conformational divergence. The goal of this study was to capture and confirm the known distinctions between *wild-type* and mutant coarse-grained simulations at reduced resolution, demonstrating the robustness of NetMD, even with simplified contact definitions.

As a final proof of concept, we applied NetMD to comparative binding simulations of GLUT1 with two chemically distinct inhibitors, *cytochalasin B* and *phenylalanine amide*, where the trajectories comprised an initial variable-length SuMD phase followed by a fixed-length GaMD phase, to test the capacity of the method for aligning asynchronous time series and resolving ligand-specific conformational signatures. This latter aspect is particularly relevant for guiding the rational design of more effective GLUT1 inhibitors, which are primary targets in many cancer treatments.

Before embedding, we applied the Shannon–entropy filter separately to each inhibitor ensemble. This step pruned the contacts showing minimal variability across all replicates for a given ligand, thereby emphasizing the most characteristic interactions of each binding pathway. Each simulation concatenated a variable-length, 1,000-to-5,200 frames, supervised MD (SuMD) simulation to a uniform-length, 2,500 frames, Gaussian accelerated MD (GaMD) simulation. This variability complicates the alignment and synchronization of time-series embeddings, which typically require comparable temporal scales to define a meaningful consensus trajectory. This was evident when mapping the DTW path on pairwise distance heatmaps, where the shortest trajectory was adaptively stretched, effectively “waiting” in embedding space until longer trajectories converged on the same conformational state. To mitigate this, we performed separate analyses of SuMD and GaMD segments.

The SuMD segments were successfully warped, revealing a shared early binding trajectory and diverging intermediate states for the two inhibitors. Notably, the alignment reconverged toward the end of the SuMD phase, demonstrating that all replicates ultimately sampled the same key binding intermediates despite asynchronous sampling. In contrast, the GaMD segments were tightly aligned, with minimal adjustments (**Supplementary Data 2a**). This consistency across replicates underscores that timing, rather than conformational heterogeneity, drives the initial alignment challenges. The alignment was quantified by computing the full pairwise normalized DTW distance matrix for all six replicates. Within‐inhibitor distances for cytochalasin B ranged from 1.87 to 2.50 (mean≈2.13), and for phenylalanine amide from 1.71 to 2.37 (mean ≈ 2.14). Cross‐inhibitor distances averaged ~ 2.36, substantially higher than either within‐group mean, highlighting the clear dynamic separation between the two binding pathways. Hierarchical clustering separated the inhibitors into distinct branches, thereby distinguishing the ligand-specific conformational pathways (**Supplementary Data 2b**).

Across all tested systems, ranging from small transporters to large respiratory complexes and ligand–protein interactions, NetMD consistently aligned and clustered trajectories without prior labeling, resolved key conformational states, and mapped divergence windows to events of mechanistic significance. This method accommodated both atomistic and coarse-grained models, variable trajectory lengths, and heterogeneous sampling protocols. These patterns, which are consistent across methods and molecular systems, underscore four key aspects: scalability, interpretability, integration potential, and limitations, which are explored in the following discussion.

## DISCUSSION

The results demonstrate that NetMD can uncover shared dynamic pathways and precisely localize divergence events across diverse molecular systems, linking algorithmic patterns directly to mechanistic insights. While our case studies focused largely on biomolecular contexts, the methodology is inherently generalizable. Any system whose evolution can be captured as a time series of interaction networks, from catalytic reaction intermediates in chemistry to defect propagation in crystalline solids, can, in principle, be analyzed using the same entropy-filtered graph embedding and time-warping framework. This flexibility arises from NetMD’s minimal reliance on system-specific prior knowledge, requiring only a consistent definition of contacts or interactions.

The performance of NetMD across atomistic, coarse-grained, classical, and enhanced-sampling trajectories shows its capacity to handle datasets that vary widely in resolution, size, and duration. In biological applications, this enables the alignment of membrane transport cycles, globular enzyme conformational changes, and ligand-binding pathways. In chemistry, it could align reactive trajectories to identify conserved transition-state geometries or distinguish solvent-dependent mechanistic routes. In materials science, it could synchronize simulations of mechanical deformation, thermal annealing, or phase transitions to detect reproducible transformation sequences and isolate the effect of compositional variations. By requiring only residue–contact graphs as input, the method remains agnostic to the simulation protocol, allowing its application to targeted simulations (e.g., SuMD), long non-targeted runs (e.g., GaMD), TMD, coarse-grained MD, or hybrid strategies without parameter reconfiguration.

In each case study, NetMD not only synchronized trajectories but also produced interpretable divergence maps. For GLUT1 and KDM6A, the divergence windows coincided with key conformational transitions and contact rearrangements linked to functional impairment in the mutants. In Complex I, increasing mutation burden correlated with greater dispersion in embedding space and larger cross-group DTW distances, consistent with experimental knowledge of functional severity. For GLUT1 inhibitors, NetMD resolved ligand-specific conformational signatures, even with asynchronous binding phases. For chemical and materials systems, analogous divergence points may correspond to bond rearrangements, nucleation events, or defect migration steps. The method’s unsupervised nature ensures that these events are detected without predefining reaction coordinates or state labels, supporting both hypothesis generation and validation.

Because NetMD operates independently of atomic coordinates once contact graphs are built, it can be readily integrated into existing MD analysis pipelines or applied retrospectively to archived datasets. For instance, studies of genotype–phenotype correlations in rare diseases could benefit from NetMD’s ability to align and compare variant versus *wild-type* trajectories, as in investigations of C5orf42 in oral–facial–digital syndrome type VI^18^ or the characterization of RASopathy-associated variants in prenatal diagnostics^19^. Similarly, NetMD could refine the interpretation of miRNA–protein interaction dynamics in cancer progression^20^, reveal mechanistic disruptions in mitochondrial respiratory complexes^21^, or dissect mutation-driven contact rearrangements in diverse pathologies^22–25^. Applications extend to meta-analyses of large-scale MD repositories, including regulatory element databases^26^ and methodological frameworks for genome research^27^, where synchronized trajectories can uncover conserved dynamic motifs across proteins, pathways, or even species.

Although NetMD scales to long trajectories and large systems, computational demands grow with the number of frames and complexity of the Weisfeiler–Lehman graph vocabulary. The embedding time increases with the number of graphs and iterations, whereas the DTW alignment cost increases with sequence length and number of replicas. Parameter choices such as Shannon–entropy cutoffs, number of WL iterations, and embedding dimensionality balance resolution against runtime. Higher cutoffs and fewer iterations accelerate computation but may omit infrequent interactions; lower cutoffs and more iterations capture finer variations but increase the cost. The availability of a flexible API with multicore parallelization mitigates these trade-offs, enabling users to tailor performance to system complexity.

The versatility of NetMD suggests that it can serve as a unifying framework for comparative MD analysis across biology, chemistry, and materials science. By enabling an unsupervised, time-resolved comparison of complex molecular systems, it opens the door to systematic mining of dynamic signatures in systems as varied as enzymes, catalysts, nanomaterials, and functional polymers. As molecular simulations continue to expand in scope and resolution, such integrative approaches will be pivotal for translating dynamic patterns into mechanistic understanding, accelerating discovery, and informing predictive models across disciplines. NetMD is freely available at https://github.com/mazzalab/NetMD.

## METHODS

### Molecular Dynamics Simulation

In this study, we exploited four different simulation strategies that span the classical and enhanced sampling MD methods. This has been demonstrated in real molecular systems.

### GLUT1 conformational transition during glucose translocation

Supervised Molecular Dynamics (SuMD)^28^ is an adaptive sampling technique that enables rapid exploration of ligand-receptor recognition pathways on a significantly reduced timescale compared to standard MD approaches. Targeted molecular dynamics (TMD)^29^ approaches are frequently used in the MD field to induce conformational changes in a target structure using steering forces, overcoming energy barriers, and allowing a more efficient conformational space exploration. Here, we extensively simulated the entire glucose pathway through its main transporter, GLUT1, in *wild-type* and mutant contexts, using a hybrid MD strategy. First, we implemented SuMD to quickly guide glucose into its central binding pocket, followed by TMD to simulate the conformational switching in the presence of glucose.

The initial outward-open conformation, together with the outward-occluded conformation, was modelled by homology modeling using the X-ray outward-open and outward-occluded conformations of GLUT3 as a template (respectively, PDB id: 4ZWB and 4ZW9), and then embedded into a lipid bilayer composed of 1-palmitoyl-2-oleoyl-glycero-3-phosphocholine (POPC) using CHARMM-GUI^30^ to simulate an accurate cellular environment *in-silico*. Conversely, the crystal structure of the GLUT1 inward-open configuration (PDB ID: 4PYP) was used as the final target for TMD simulations. Single point mutations Arg333Trp (R333W) and Arg333Gln (R333Q) were introduced using ChimeraX^31^. Both *wild-type* and mutant systems were inserted into a simulation box filled with TIP3P and Na+/Cl− counter ions to neutralize the overall charge. Finally, a D-glucose molecule was inserted into the simulation box far from the GLUT1 central cavity and slowly guided into the X-ray-determined binding pocket, while exploring a variety of potential binding pathways.

The obtained systems were energy-minimized by applying the steepest descent method, followed by the conjugate gradient method, and then gradually heated and equilibrated for 10 ns using a time step of 1 fs. Ten replicates of each simulation were conducted using the Amber ff14SB force field. The SuMD phase consisted of short consecutive simulations of 300 ps while the TMD phase was divided into two different run of 10 ns, the first with geometrical constraint that guided the outward-open→outward-occluded transition, while in second run we applied steering forces to drive both the outward-occluded→inward-open transition and the glucose along the Z-axis towards the cytoplasm. All simulations were performed using Amber 22^32^.

#### 1.2 Gaussian accelerated MD of KDM6A interaction with H3 histone

Gaussian-accelerated Molecular Dynamics (GaMD)^33^ represents one of the most robust MD approaches to simulate protein conformational transitions among many biological processes. By adding a harmonic boost potential that follows a Gaussian distribution, GaMD expands sampling by orders of magnitude without the need for predefined reaction coordinates.

Here, we simulated the impact of a known pathogenic missense mutation, Arg1255Trp (R1255W), on KDM6A-H3 interaction. Specifically, the catalytic Jumonji (JmjC) domain of KDM6A, a histone demethylase, was simulated in complex with the H3 histone using GaMD, following the system preparation and simulation protocol illustrated in^15^. In brief, after standard minimization and equilibration steps, as described above, the *wild-type* and mutant systems were simulated five times for 250 ns each. The Amber ff14SB force field was used to parameterize the amino acids, whereas the Zinc AMBER force field (ZAFF) was employed for the Zn(II) ion. All simulations were performed using Amber 22^32^.

#### 1.3 Coarse-grained simulations of single and triple mutant mitochondrial CI complex

Coarse-grained (CG) molecular dynamics simulations^34^ are widely used to simulate biomolecular systems on large time and size scales. In a CG simulation, groups of atoms are represented as a single particle, reducing the computational complexity while retaining the essential structural and dynamic properties of a biomolecule.

Here, we tested our NetMD approach on the CG-MD simulations described in^16^ and made available at Zenodo (https://doi.org/10.5281/zenodo.10355270). In this study, we employed a CG representation of mammalian respiratory complex I (CI) to determine the impact of putatively pathogenic mtDNA variants, alone or in combination, on the conformational transition of the transmembrane helix (TMH3) of MT-ND6, which carries the m.14484T>C primary mutation and is fundamental for CI function. We selected only human-open systems for the *wild-type*, single mutant, and triple mutant.

### Comparison of the binding mechanism of GLUT1 inhibitors employing Su-GaMD

Rational integration of different MD sampling techniques is fundamental for the accurate characterization of complex biological events, such as ligand-induced activation/inhibition processes. Su-GaMD^35^ is an enhanced sampling technique that incorporates the SuMD and GaMD approaches described in previous sections, providing a powerful method to simulate the entire ligand recognition process together with the occurring receptor conformational changes, enabling a deeper understanding of the ligand mechanism of action.

Here, we compared the full binding mechanism of two known GLUT1 transport inhibitors, cytochalasin B and phenylalanine amide, whose complex structures with the transporter are available in the PDB (5EQI and 5EQG, respectively), using the Su-GaMD protocol described in^35^. GLUT1 was inserted into a POPC bilayer, and each inhibitor was initially placed approximately 50 Å away from the respective binding pocket. GaMD acceleration parameters were derived from prior GaMD simulations. Short 600 ps trajectories were employed for the SuMD steps, followed by 100 ns of GaMD. Thus, following the standard minimization and equilibration protocol described in Section 1.1, each system was simulated in triplicate using the Amber 14SB force field. All simulations were performed using Amber 22^32^.

### MD trajectories processing

#### MD trajectory to Graph conversion

From the raw molecular dynamics (MD) simulation trajectories, we derived time-resolved contact maps by generating edge lists that captured all residue–residue interactions across frames. Trajectories were processed using GetContacts (https://getcontacts.github.io/), which enabled the systematic extraction of pairwise residue interactions on a per-frame basis. For each frame, we recorded a list of interacting residue pairs, annotated with the corresponding frame index. This approach yielded a temporally indexed contact graph for each replica, an efficient yet expressive representation that preserved the full connectivity of the system while maintaining both spatial and temporal resolution. These contact graphs served as the foundation for the subsequent frame-by-frame analysis of interaction dynamics across replicas.

For the coarse-grained system, we first identified all residues that fell within a 15 Å radius of the mutation site in at least one frame. This spatially filtered node set was then used to build per‐frame edge lists for each replica. At every time point, an undirected edge was placed between any two of the selected residues whenever their pairwise distance dropped below 6 Å.

#### Graph edge pruning

The residue-contact graph was pruned using an entropy-based metric to enhance the representativeness of the dataset. For all frames of all simulation replicas of a molecular system, we calculated the Shannon entropy, a widely recognized metric for quantifying variability and disorder in biological systems. In this application, entropy provides an *ad hoc* method for capturing the dynamism of residue interactions, which is essential for understanding conformational flexibility and functional hotspots^36^. It then provides a quantitative measure of how interactions fluctuate over time.

This filter excludes (i) low-frequency noisy contacts and (ii) high-frequency invariant contacts that, while structurally stabilizing, provide little discriminative value between frames. The choice of a frequency threshold is critical because it aims to emphasize case-specific interaction patterns. Lower thresholds may capture more interactions, potentially increasing sensitivity to subtle variations. Higher thresholds provide stricter criteria and focus on only the most dynamic interactions. Nevertheless, the possibility of tuning this value ensures that the method can be tailored to specific biological questions or data characteristics.

After applying the entropy filter, we extracted subgraphs that contained only the retained edges and their incident nodes. In such graphs, nodes represent residues and edges denote the informative contacts between residues. Nodes were annotated using multiple features. We specifically used the position of the residue within the amino acid sequence as one such feature, which inherently captures information regarding both the primary and tertiary structures of the protein. This ensures that no information about the residue identity is lost during graph construction, thus maintaining a direct link to the biological entities of the molecule. Importantly, the number of nodes remained consistent across all graphs because they represent the same set of residues in the molecular system. This ensures that variations in graph properties are strictly related to changes in residue interactions over time and allows us to accurately capture the system’s behavior over time and across different replicas, providing residue-level insights^37^.

#### Graph Embedding

Embeddings were generated from the graphs using the Graph2Vec^12^ method implemented in the KARATECLUB^38^ Python library. Graph2Vec is a graph-embedding method inspired by the document-embedding approach Doc2Vec^39^. It represents entire graphs as fixed-size vectors by learning a continuous vector space where structurally similar graphs are mapped closer together. Graph2Vec employs the Weisfeiler–Lehman (WL) kernel to summarize the graph topological patterns. At different scales, each node’s representation is updated by combining it with that of its immediate neighbors. Thus, after several rounds, the final representation progressively reflects larger neighborhoods (i.e., both direct and indirect contacts). This makes the WL kernel particularly well suited for molecular graphs, where the complex connectivity of residues must be precisely represented, and by the end of the process, the refined node representation encodes both local and intermediate structural features of the graph. After applying the WL kernel, the Graph2Vec algorithm partitions the graph into a collection of rooted subgraphs (*bag-of-subgraphs*), each of which corresponds to a node and its local neighborhood defined by the previous step. These subgraphs are the key elements for feature extraction, and a feature vector is constructed for each graph by counting the occurrences of different subgraph patterns and generating a distribution of subgraph labels.

Then, a skip-gram model is trained like the approach used in Word2Vec^40^, which learns to represent graphs as real-valued vectors by analyzing their surrounding context (*bag-of-subgraphs)*, thereby capturing the relationships between them, like how a language model learns word embeddings based on their context within sentences. Using this approach, Graph2vec can learn low-dimensional embeddings that encode the relationships and co-occurrence patterns of subgraph labels that reflect the topological properties of the graph.

In our workflow, we selected the following parameters for the Graph2Vec method: (i) the dimensionality of the embeddings was set to 16, balancing the need for a compact representation with the ability to capture meaningful structural patterns; and (ii) the number of WL iterations (*wl_iterations*) was set to three. This allows the model to incorporate subgraph information up to three hops away from each node, thereby capturing both local and intermediate graph structures. Additionally, the downsampling rate was fixed at zero, ensuring that no subgraph patterns were discarded during the embedding process. All other parameters were kept at their default values as they adequately met the needs of our analyses without requiring additional adjustments.

The embedding result was a three-dimensional matrix of size *NxRxD*, where *N* corresponds to the number of frames, *R* to the number of replicas, and *D* to the embedding dimensions: more specifically, each graph, representing a single frame from a specific replica, was encoded as a vector of size *D*.

To facilitate the visualization of the embedding matrix, Principal Component Analysis (PCA) was used to retain 90% of the variance, reducing the dimensionality while preserving the most significant features of the data. This step ensures that the most important patterns in the embeddings are maintained, thereby allowing for a meaningful visual representation. Subsequently, Spectral Embedding was used to project the data into a low-dimensional space based on the connectivity structure of the graph. This two-step approach, PCA followed by Spectral Embedding, enhances the visualization of relationships and clustering among frames and replicas within the embedding space. Moreover, the analysis provided hints on how graphs group according to their structural similarities, providing a more intuitive interpretation of the data, and helping in the identification of distinct patterns or clusters.

#### Consensus Identification via Dynamic Time Warping

We considered the set of embedding vectors from each replica as a time series of size *NxD*. Each embedding vector of dimension *D* encodes a snapshot of the conformation of the protein at a given frame in *N*, faithfully describing its dynamic behavior throughout the simulation. Consequently, to capture a consensus representation of the embeddings across all replicas, we employed the Dynamic Time Warping Barycenter Averaging (DBA) technique ^13,41^.

DBA is a method for computing the barycenter of multiple time series while accounting for their temporal alignment, thereby providing a central representation of the data. It aims to identify a central trajectory that minimizes the average distance between time series while also considering their nonlinear temporal shifts, making it particularly suitable for data with varying lengths and misalignments. It is based on Dynamic Time Warping (DTW)^42^, which allows the comparison of time series with different lengths or temporal shifts. DTW identifies an optimal alignment by stretching or compressing segments of the time series to minimize the overall discrepancy between them. This alignment is crucial for capturing the true temporal dynamics of the system, even in the presence of distortions in the timing of individual time series.

Through this approach, we obtained a “*consensus embedding*,” which captures a generalized, stable pattern of the protein’s dynamics across all replicas. However, in addition to computing the barycenter, we aligned all replicas to the consensus embedding. This step allowed us to normalize the dynamic behavior of each replica relative to the central representation, ensuring that all replicas were compared in a consistent temporal frame. To measure the dissimilarities between each time series, we introduced a scoring function based on the normalized DTW distance. This function accounts for differences in the lengths and temporal shifts of the time series. The normalized DTW formula is as follows:

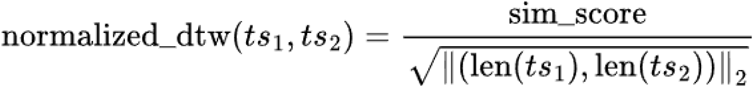

where *sim_score* is the similarity score obtained from the DTW path computation between time series *ts1* and *ts2* and the denominator normalizes the score by considering the lengths of the time series.

#### Data-Driven Cluster Detection and Outlier Removal

To explore the relationships between replicas, we performed hierarchical clustering based on the pairwise distances computed between each time series. A linkage matrix was computed using Ward’s method to minimize the total within-cluster variance.

To identify the optimal number of time-series clusters, that is, the number of distinct groups exhibiting similar temporal patterns, we devised two complementary, data-driven strategies that do not require prior specification of the number of clusters. (i) The *largest gap method* detects the point where a substantial increase in linkage distances occurs, assuming that it represents a natural division between clusters. The reasoning behind this is that stepwise dendrograms frequently create higher jumps in linkage distances whenever its elements lack clear cluster affinity, which can serve as indicators for identifying the optimal number of clusters^43^. (ii) The classical *elbow method* is based on the principle of diminishing returns in the cluster cohesion. It assesses the linkage distances between merges in the hierarchical clustering process and identifies the point at which adding more clusters no longer significantly reduces within-cluster variance. This point, known as the “elbow,” marks the optimal balance between excessively fragmented data into too many clusters (oversegmentation) and grouping dissimilar elements into too few clusters (undersegmentation). By detecting the steepest decline in distances between successive merges, the method estimates the dendrogram’s inflection point, enabling data-driven determination of optimal cluster partitioning.

Finally, we implemented an iterative pruning method to further refine the reference embedding. The main goal of this approach is to identify and eliminate replicas that deviate most significantly from the consensus, thereby ensuring that the final central representation robustly reflects the core dynamics. At each step, the barycenter was computed and the replica with the largest deviation was removed until only the two closest replicas (both at the same distance from the barycenter) remained. This process allowed us to progressively eliminate outliers and minimize the influence of extreme or atypical conformational states.

#### Change-Point Detection via Temporal Deviation Profiling

To detect anomalous dynamic behaviors within individual replicas, we computed the pairwise Euclidean distance at each aligned time step (frame), comparing each replica to a reference trajectory selected as the closest to the consensus. This procedure generates a temporal deviation profile that captures the localized structural divergence over time.

To reduce noise and enhance the interpretability of these temporal distance signals, we applied a centered moving average filter with an odd-sized sliding window. MD data can exhibit high-frequency noise owing to intrinsic thermal fluctuations, sensitivity to initial velocities, finite sampling, and numerical integration artifacts. With this data, a moving average can smooth out these short-term irregularities.

We then used the Pruned Exact Linear Time (PELT) algorithm, an efficient change-point detection algorithm designed to identify points in a time series in which the underlying statistical properties change. PELT aims to determine a set of change points 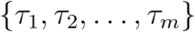 that minimize the following objective function:

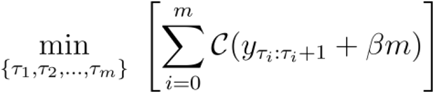

where i) *C* is the cost of each segment; ii) *β* is the penalty term; and iii) *m* is the number of change points. More specifically, we opted for a linear cost function that assumes that, within each segment, the observed signal can be described by a straight line (constant slope and intercept), while also penalizing deviations from that line, thereby allowing the algorithm to identify points at which the topology of the system changes significantly. Here, a “segment” is simply the interval between two successive change points, over which the deviation profile is treated as stationary and can be changed to tune the sensitivity accordingly: smaller values yield shorter and thus more segments by checking for mean shifts every few frames, whereas larger values produce fewer, longer segments. The penalty was logarithmically scaled with the number of frames, allowing for a consistent performance across datasets of varying sizes.

## FIGURES

**Supplementary Data 1.**
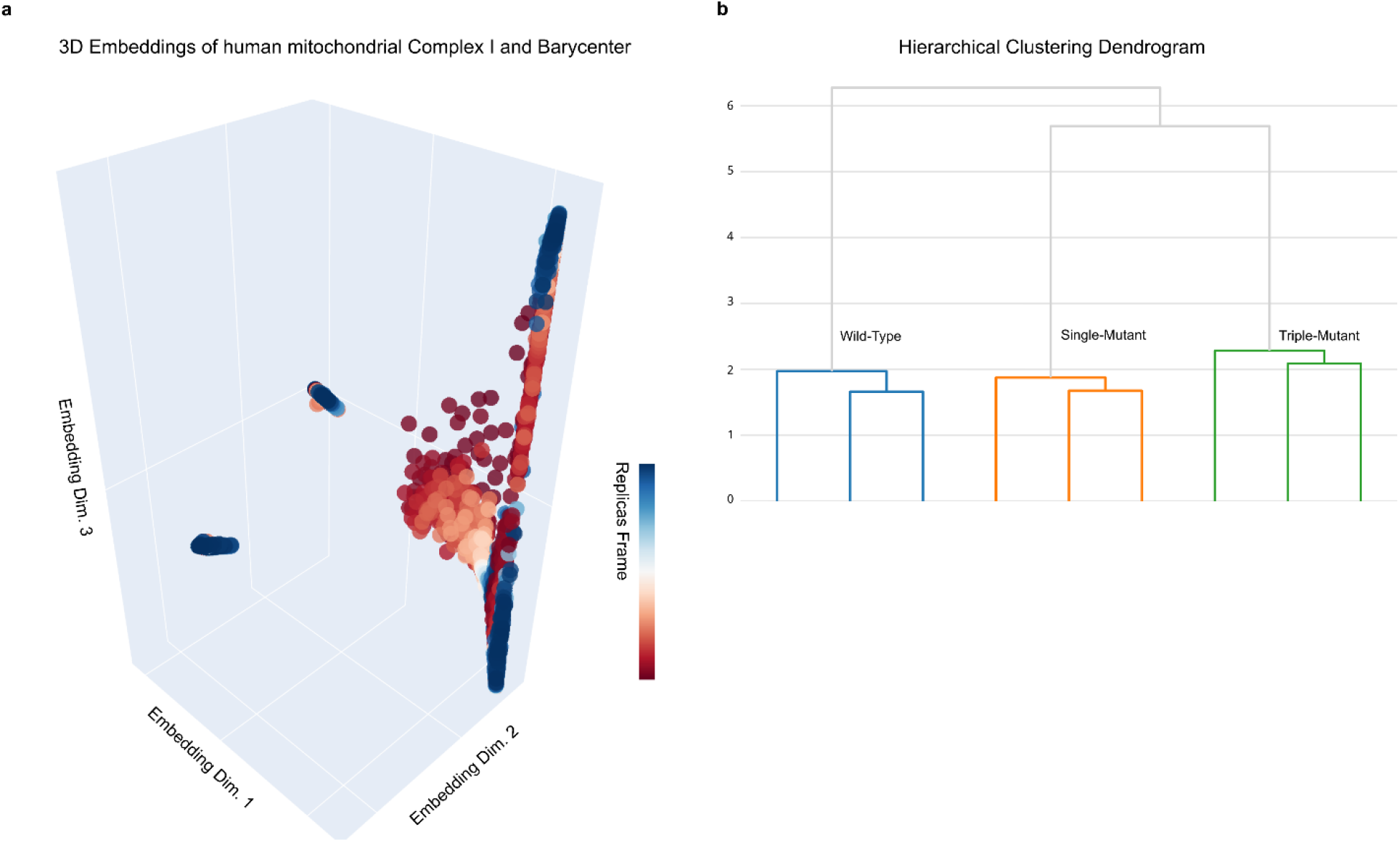
(a) Combined 3D embeddings of coarse-grained simulations of the *wild‐type*, single mutant, and triple mutant. (b) Final hierarchical clustering dendrogram computed on the combined embeddings, illustrating the three major branches corresponding to the *wild-type* and the mutants; in particular the triple mutant system is the farthest from the *wild-type*.

**Supplementary Data 2.**
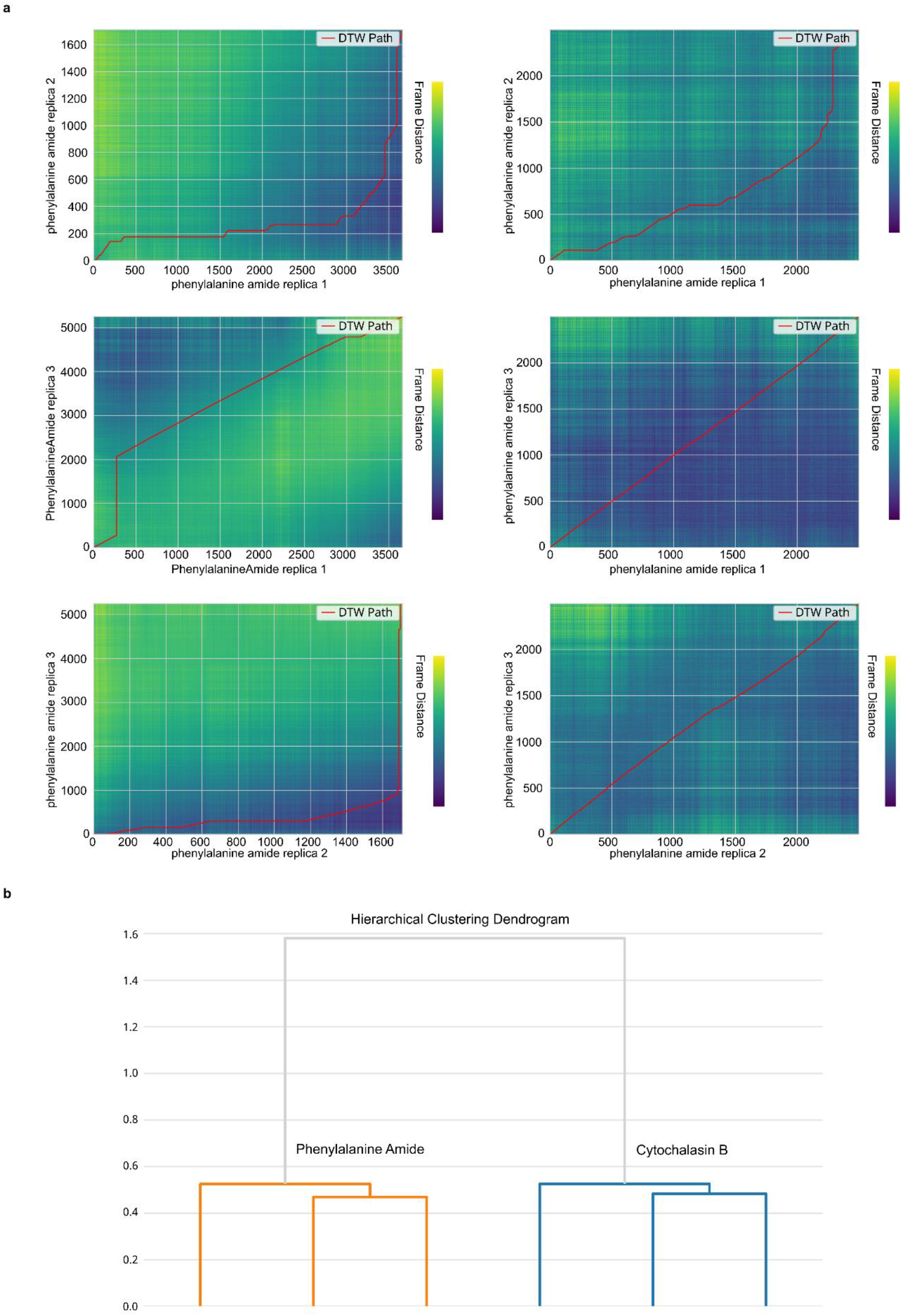
(a) Dynamic time–warping (DTW) alignments of ligand trajectories from SuMD (left column) and GaMD (right column) simulations. Each heatmap shows the pairwise frame distances between three independent replicates (rep1 vs. rep2, rep1 vs. rep3, and rep2 vs. rep3) of the phenylalanine amide system, with the optimal DTW path overlaid in red. In SuMD, unequal trajectory lengths produce pronounced deviations of the red path from the diagonal (time warping), whereas in GaMD (in which all trajectories have the same length), the DTW path remains nearly linear. (b) Hierarchical clustering dendrogram: All SuMD+GaMD replicates from the phenylalanine amide system (orange) form one cluster, whereas those from cytochalasin B (blue) form the other, illustrating that the learned embedding robustly discriminates between the two distinct inhibitor–protein systems.

**Supplementary Table 1**. Contact frequencies for *wild-type* and R333Q over a fixed time window (frames 205–432) and their WT/MUT odds ratios: values higher than 1 indicated contacts favored in *wild-type* and values lower than 1 indicated contacts more conserved in the mutant.

**Supplementary Table 2**. Contact frequencies for *wild-type* and mutant over a fixed time window (frames 1015–2202) and their WT/MUT odds ratios: values higher than 1 indicated contacts favored in *wild-type* and values lower than 1 indicated contacts more conserved in the mutant.

## Data availability

The residue-residue contacts files extracted from the MD trajectories of the GLUT1, KDM6A, and GLUT1-inhibitors systems that were analyzed in this study are available at ^17^. The landing page of the GitHub repository associated with this paper is https://github.com/mazzalab/NetMD. The documentation is available at https://mazzalab.github.io/NetMD.

## Code availability

Code to reproduce the analyses performed in this study is provided as a Jupyter Notebook at https://github.com/mazzalab/NetMD (**tutorial** folder).

## Acknowledgements

This project was funded by the Italian Ministry of Health (RC2025). We additionally acknowledge ISCRA (IscrB_GLUT1-MD - HP10BC3V4L) for awarding this project access to the LEONARDO supercomputer owned by the EuroHPC Joint Undertaking, hosted by CINECA (Italy).

