## Supplementary figures and images for "NetMD: Unsupervised Synchronization of Molecular Dynamics Trajectories via Graph Embedding and Time Warping"

### Supplementary Data 1

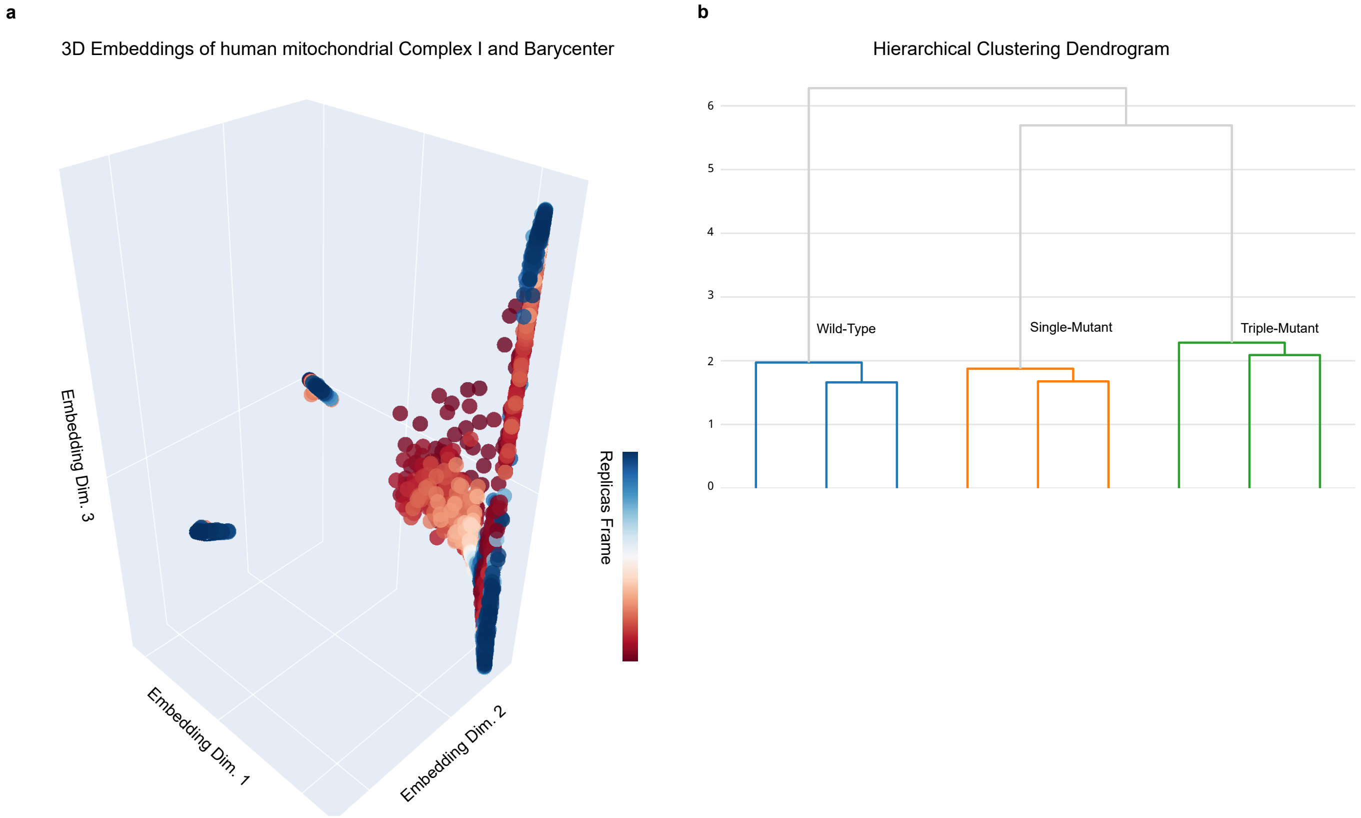

### Supplementary Data 2

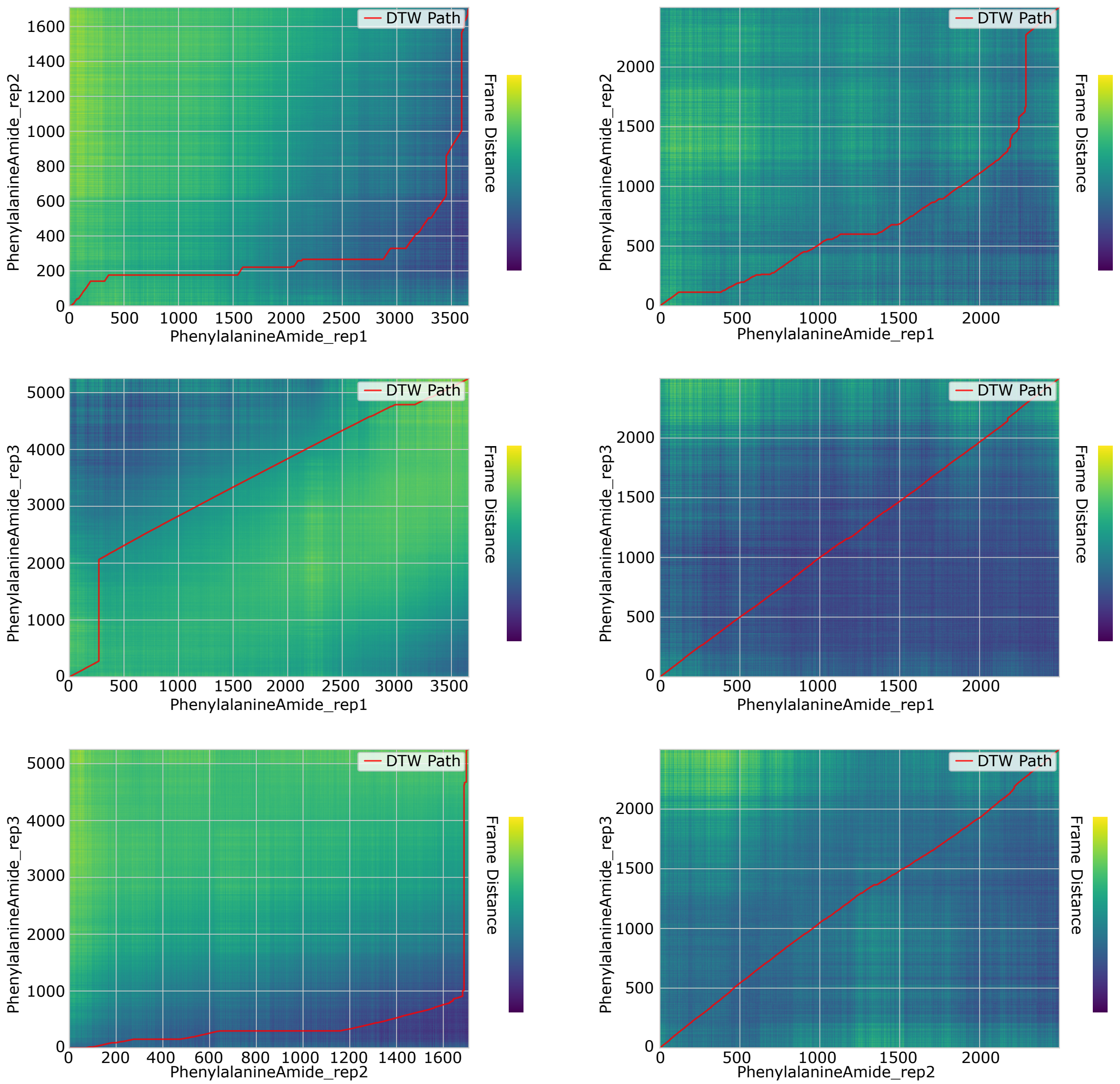

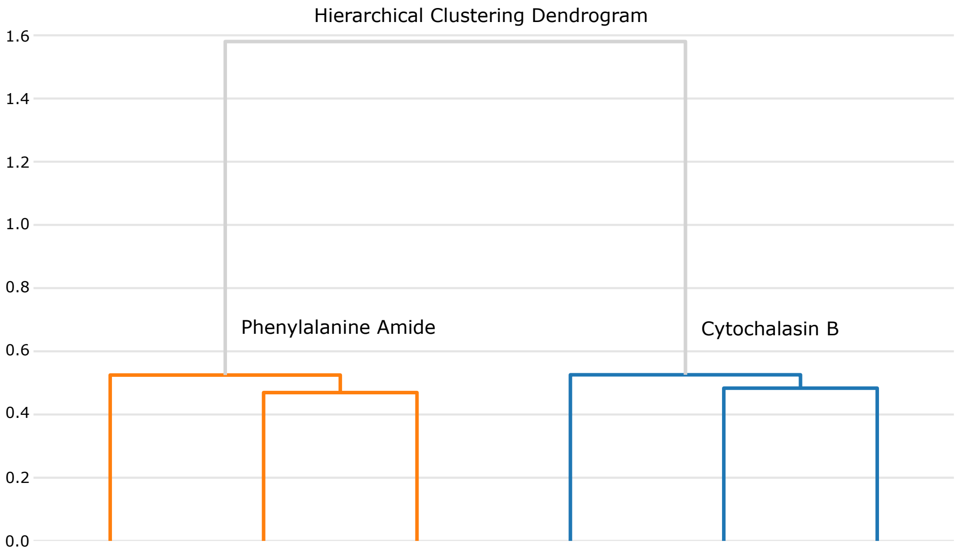
